# High-throughput, low-cost and rapid DNA sequencing using surface-coating techniques

**DOI:** 10.1101/2020.12.10.418962

**Authors:** Yanzhe Qin, Stephan Koehler, Shengming Zhao, Ruibin Mai, Zhuo Liu, Hao Lu, Chengmei Xing

**Affiliations:** MGI, BGI-Shenzhen, Shenzhen 518083, China; Harvard T.H. Chan School of Public Health, Boston, MA 02115, USA; BGI-Shenzhen, Shenzhen 518083, China

## Abstract

The speed^1–3^, expense^1–4^ and throughput^2^ of genomic sequencing impose limitations on its use for time-sensitive acute cases, such as rare^4,5^ or antibiotic resistant infections^6^, and large-scale testing that is necessary for containing COVID-19 outbreaks using source-tracing^7–9^. The major bottleneck for increasing the bandwidth and decreasing operating costs of next-generation sequencers (NGS) is the flow cell that supplies reagents for the biochemical processes; this subsystem has not significantly improved since 2005^10–12^. Here we report a new method for sourcing reagents based on surface coating technology (SCT): the DNA adhered onto the biochip is directly contacted by a reagent-coated polymeric strip. Compared with flow cells the reagent layers are an order of magnitude thinner while both the reagent exchange rate and biochip area are orders of magnitude greater. These improvements drop the turn-around time from days to twelve hours and the cost for whole genome sequencing (WGS) from about $1000 to $15, as well as increase data production by several orders of magnitude. This makes NGS more affordable than many blood tests while rapidly providing detailed genomic information about microbial and viral pathogens^6,13^, cancers^14^ and genetic disorders for targeted treatments^6^ and personalized medicine^6,15^. This data can be pooled in population-wide databases for accelerated research and development as well providing detailed real-time data for tracking and containing outbreaks, such as the current COVID-19 pandemic.

## Introduction

High-throughput DNA sequencing is becoming ever more commonplace for public health^16–18^, and is a powerful tool for dealing with pandemics^19^, such as tracing the origins of SARS-CoV-2 outbreaks^7,20^ and following the virus’s evolution^6,21,22^. In China several COVID-19 outbreaks were successfully contained using source-tracing based on NGS sequencing, because their sources were viral strains that had evolved outside of China^8,9^. But on the scale of the ever-growing current pandemic that is approaching 100 million infections and millions of deaths^23^, the limited data throughput of state-of-the-art NGS (in two days a mainstream NGS platform produces 300Gb or whole genome sequences (WGS) for only 3 people^24^), slow turn-around times (test results typically take two or more days^24^), and high expense of current sequencing (about $1000 per WGS or 100Gb) limit their use for tracking and containing outbreaks^19^.

The core component of NGS is the flow cell where the bioreactions take place, and after significant improvements in 2005 the operating costs and throughput over the last five years have remained unchanged, see Fig. S1. It is a microfluidic device with a gap of ~100 μm and contains multiple lanes of attached single-stranded DNA molecules^12^ that undergo a sequence of biochemical reactions and are optically imaged for base calling after each cycle^25^. Recent attempts at improving throughput include reducing the spacing between DNA attachment points^12^; however, this reduction is curtailed by the resolution limits of optical imaging^26^. Another approach for improving throughput and turn-around times is increasing the size of the flow cell^27^ and boosting its flow rate. But there are limits to the flow cell’s structure: larger flow cells have a propensity for non-uniform reagent flows, see Fig. S2; higher flow rates, reduced gap and the concomitant cubic increase in pressure^28^ causes bending and delamination of the coverslip. The fastest NGS system can generate about 3Tb (30 WGS) per day with a typical cost of about $1000/person and turn-around time of 2 days^29^.

Here we present a surface-coating technique (SCT) that dramatically improves data throughput and reagent usage by rapidly applying a thin reagent coating on a large biochip covered with a microarray of 5×10^10^ DNA molecules. This increases the data size and exchange rates by orders of magnitude, see Fig.1. A PET polymeric strip coated with reagents directly contacts the biochip for fast sourcing of reagents to the DNA. Bands of reagents are arranged sequentially on the strip so that step-wise lateral displacements provide reagents for each step of the sequencing-by-synthesis biochemical reactions^25,30^, as shown in Fig. 2a. This sidesteps issues associated with flowing reagents through a thin flow cell, such as uneven reagent distribution and high pressures necessary to quickly exchange reagents. Our prototype biochip area is 225 cm^2^ which is two orders of magnitude larger than current single-lane flow cells and can be further increased several orders of magnitude. Moreover, the reagent layer thickness is far thinner than that of flow cells, which reduces reagent waste from 99.9% to about 99% and can possibly be reduced to about 90% (see calculation section in supplementary information); this is significant because reagents currently account for 80-90 percent of operation costs^31^. The speed of coating the biochip with reagents is 80 m/min, which is over an order of magnitude faster than that in a flow cell, see Fig. 1b, and in principle can be increased to industry levels of 300 m/min^32^. A robotic arm transfers the biochip from the prototype SCT subsystem to the commercial imaging subsystem (DNBSEQ-T10 by MGI) for base calling between biochemical reaction cycles, as shown schematically in Fig. 2b. The data quality is comparable to that of a flow cell, as shown in Fig. 3d. For 20 cycles our unfiltered Q30 exceeds 80% and other key parameters such as the lag and run on are acceptable, as shown in Figs. 3e and f.

**Fig. 1.**
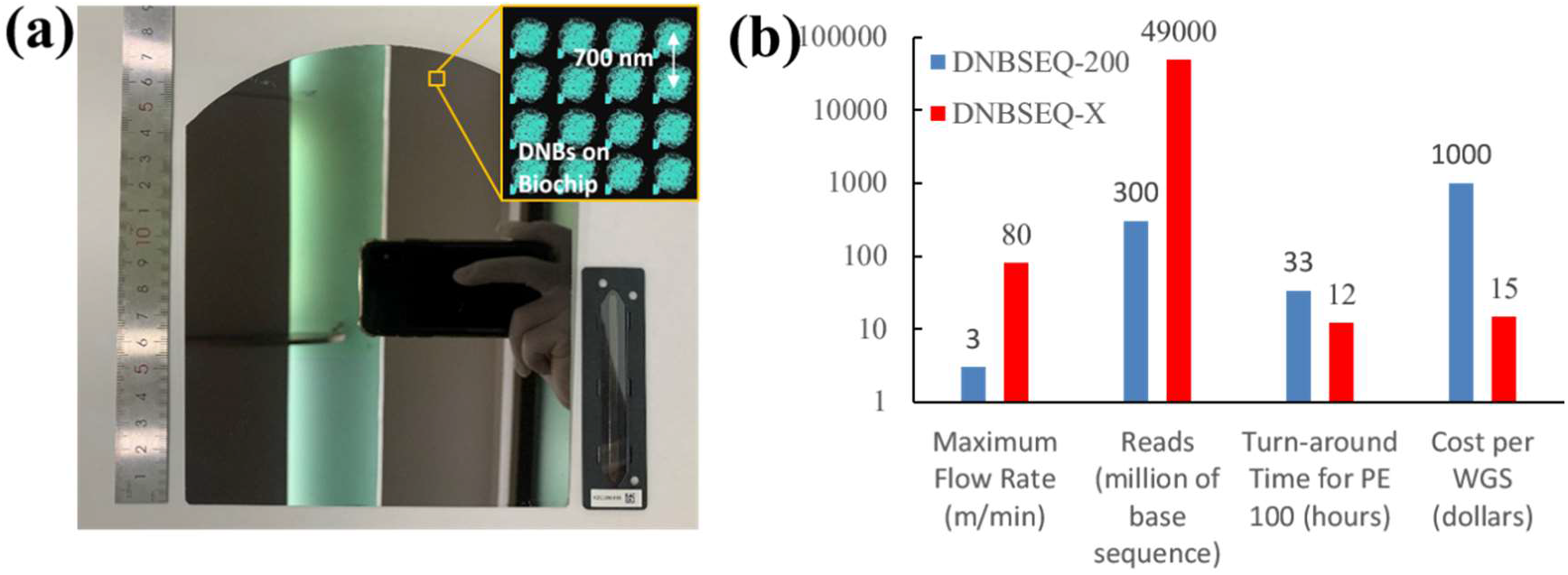
Advantages of SCT. (a) Size comparison of biochips for SCT and an NGS flow cell. Inset show pattern array loading with DNBs. (b) Comparison of the flow rate, reads, turn-around time and cost for a WGS for a conventional DNBseq-200 single-lane flow cell and out SCT prototype.

**Fig. 2.**
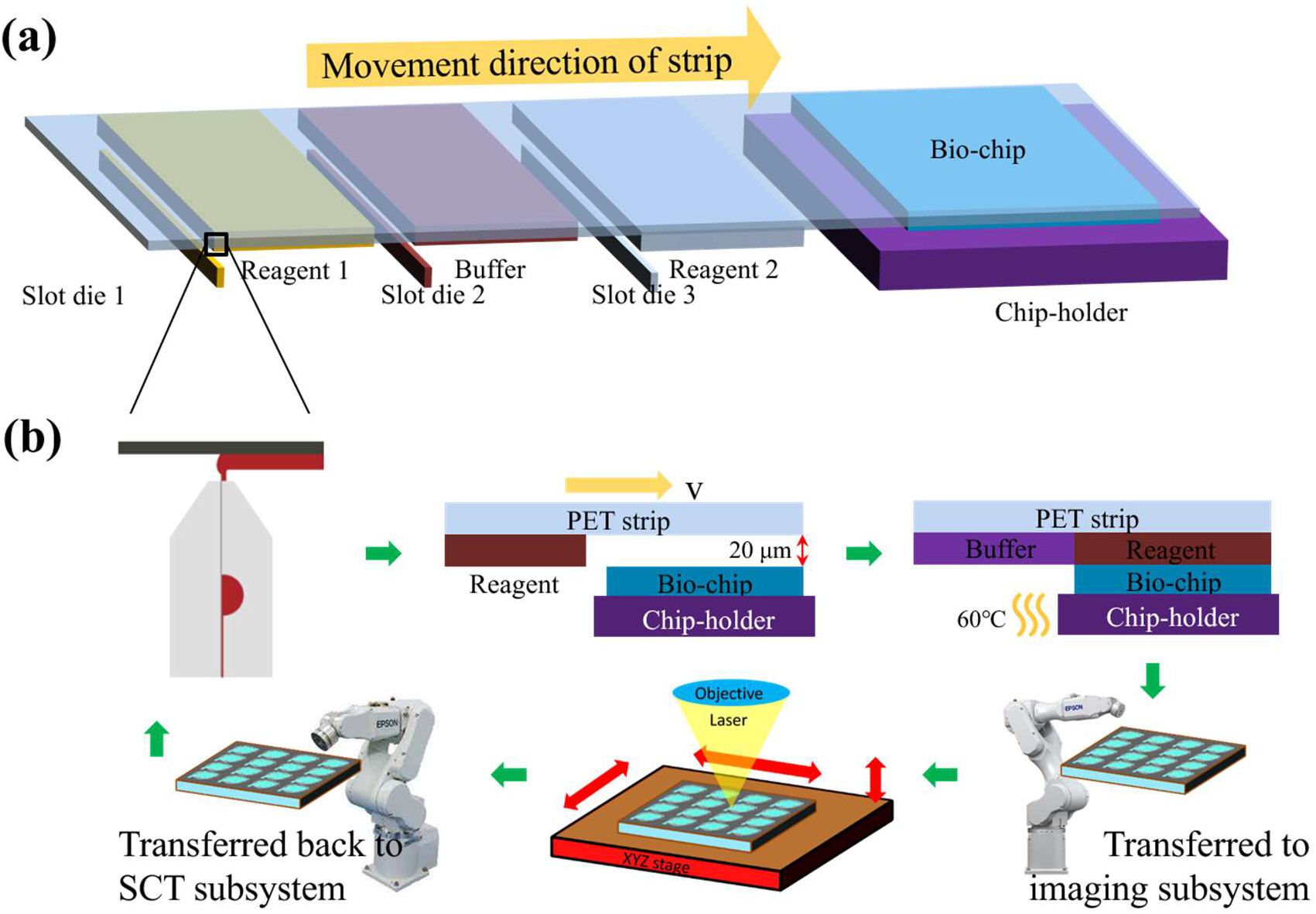
Principle and set-up of the strip-coating technique (SCT) subsystem. (a) Schematic of the coating and replacement process for sequencing reagents, using a moving hydrophilic PET strip to drive flow. (b) Schematic of the workflow for SCT NGS: coating the strip, advancing the strip to source reagents and apply heat, transfer the biochip to the imaging subsystem, and returning the biochip for supplying the next reagent.

**Fig. 3.**
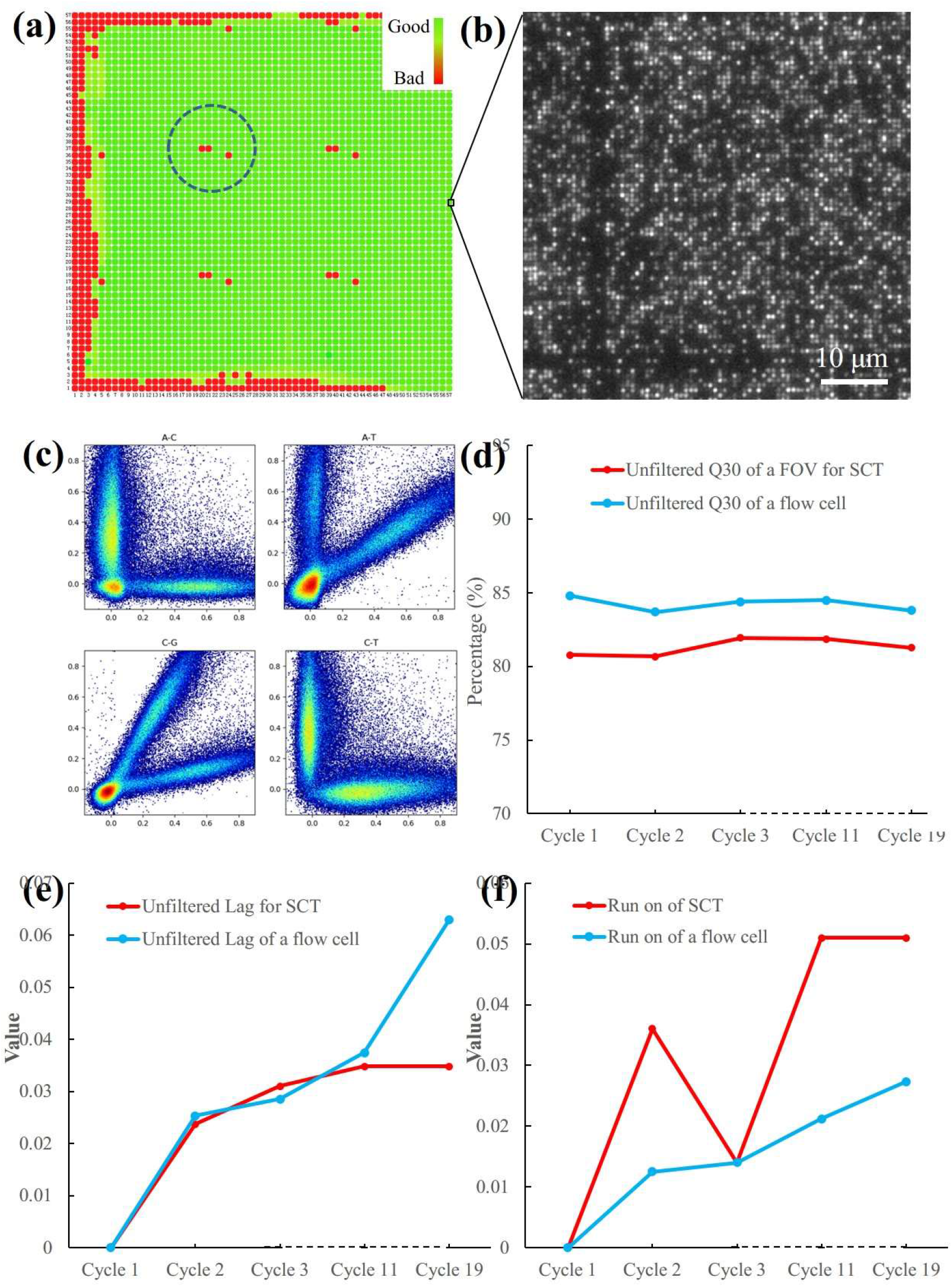
Data quality achieved with our SCT prototype. (a) Base-calling information content (BIC) heatmap for a sequencing chip. Green indicates acceptable quality for base calling. The dashed white circle indicates one of the four regions used for image registration. (b) A fluorescent image zoomed in to show a square region of micropatterned DNBs (c) Crosstalk plots for the A-C, A-T, C-G, and C-T fluorescent channels, respectively, which show good channel separation. (d) Comparison of unfiltered Q30 of a FOV and that of a flow cell for the first 19 cycles. (e) The lag and (f) run on of the first 19 cycles.

## Results and Discussion

The main challenges for replacing the NGS flow cell with SCT were achieving uniform coatings on the PET strip, maintaining the reagent-filled gap between the PET strip and the biochip, and mitigating effects of evaporation resulting from our open-cell approach. The SCT subsystem was designed to be compatible with base calling using our imaging subsystem (DNBSEQ-T10 by MGI) with minimal modifications by patterning a DNA array similar to commercial NGS platforms, and the main challenge was adapting a robotic arm for transferring the biochip between the two subsystems.

To create uniform reagent coatings on the corona-pretreated PET strip we used slot dies on an industrial roller and matched the reagents’ surface properties to that of the strip, see Fig. 2a, S3, and S4a. There are six slot dies for each of the reagents (including buffer) required for sequencing, that apply 20 um reagent layers to the strip with roller speeds of 80 m/min and motor response times of 10 ms. The biochip is placed downstream from the slot dies with a gap that matches the reagent layer thickness – see Fig. 2b. To maintain the layer’s uniformity, we placed adhesive tape of the same thickness at the edges of the biochip, see Fig. S5. Drying is not an issue because the tape slows evaporation and the NGS reaction steps take less than a minute.

The length of the moving reagent band necessary for supplying and flushing reagents scales with that of the biochip. We find reagent bands that are twice the length of the biochip replace previous liquid and effectively recoat the biochip, see Fig. S6, video clip 1 and simulation 1 in supplementary information. Directly coating the PET strip for contact with the biochip is far more efficient than using flow cells: the reagent layer is much thinner (20 μm, which in principle can be reduced to 2 μm^33^) and there is no tubing that needs to be flushed, see simulation 3.

After executing a bio-chemical reaction cycle the biochip is removed from the PET strip with a robotic arm and placed in our commercial DNBSEQ-T10 imaging subsystem for base calling, see video clip 2. As the biochip is exposed to air in this step the robotic arm must move quickly to prevent drying of the DNA. During imaging the biochip is placed under an objective immersed with an imaging solution. When the biochip is neither subject to biochemical reactions nor being imaged, we place it in a buffer bath to prevent evaporation.

We verified the accuracy of the genomic sequences from our SCT subsystem. Aside from the biochip’s edges where the presence of the tape disrupts the flow and heat transfer, the fluorescence signal was sufficient for the remaining 92% of the biochip for base-calling, see Figs. 3a and 3b. We verified that the fluorescent channels had good separation, see Fig. 3c. Similar to our commercial NGS platform^34^, our Q30 remains above 80% after 20 cycles, and both lag and run on are comparable, see Figs. 3d, e and f.

Finally, we compare our method with state-of-the-art NGS genomic sequencing in terms of turn-around time, bandwidth and cost, as shown in Fig. 1b. Since the reagent exchange is an order of magnitude faster than flow cells, PE100 or 212 cycles and reads can be completed within 12 hours (8.8 hours minimum with optimistic parameters for all subsystems) rather than two days. A single lane chip system such as DNBSEQ-200 produces 300Mb per day while our single chip system produces 20 Tb, which is due to much faster reagent exchange and larger biochip area. Behind the rollers the SCT strip has a 2m x 0.4m region available for coating biochips, which is sufficient for 266 of our 225 CM^2^ biochips that would generate 8500Tb per day. Since the data production of a single imaging subsystems is 20Tb per day, 425 such subsystems running in parallel would be required, as shown in video clip 2. SCT has a much thinner reagent layer and unlike flow chips and has no tubing which decreases reagent usage by orders of magnitude. Since the reagent cost accounts for 80-90% of current NGS platforms, this decrease in reagent usage of SCT drops the operating costs from about $1000 to about $15 for WGS.

## Conclusions

We developed a new NGS subsystem that replaces the flow cell with SCT where the throughput is increased by two orders of magnitude and the turn-around time has decreased from several days to 12 hours without compromising data integrity. Moreover, much less reagents are wasted using SCT, thereby reducing costs by almost two orders of magnitude. We anticipate that significant improvements to our SCT prototype can be made. If adopted, our technology would rival standard diagnostics, such as blood tests and biopsies, in terms of cost and speed while providing vastly more detailed and comprehensive information in terms of the whole genome that can be collected in massive population-wide databases^**35**^. This in turn will democratize personalized medicine, dramatically accelerate the pace of research and development of therapies, and provide epidemiologists with detailed data for combating outbreaks, such as those of the current COVID-19 pandemic.

## Materials and Methods

Biochips were fabricated from 8-inch silicon wafers and handles were glued on for robotic handling. Deep ultraviolet (DUV) lithography was used to fabricate a 15 x 15 cm array of circular wells with diameter of ~250 nm with center-to-center separation of 700 nm. DNA nanoballs (DNBs) were extracted from E. coli ATCC 8739 through fragmentation and recursive cutting with type IIS restriction enzymes, directional adapter insertion, and replication with Phi29 polymerase^30^. Each well was loaded with one DNB using loading and post-loading procedures, see ref 30 for details. To avoid drying the DNB-loaded biochips were stored in buffer.

We purchased corona - pretreated poly (ethylene terephthalate) (PET) rolls of from Shanghai Fuzhong Limited that were 100 kilograms of transparent, 100um thick membrane with custom cut width. The roll we use is 15 cm in radius which contains more than 1000 meters membrane and supports up to 300 cycles. Our industrial roll coater has speeds that are adjustable up to 200 m/sec with response times of 10 ms. It is equipped with 6 slot dies made by Foshan edge development mold mechanical technology Inc. The strips were under 20 kg tension and the slot dies were adjusted to create 20 μm thick layers at speeds ranging from 0.3 m/min to 80 m/min.

In order to achieve uniform coatings, we matched the reagents’ chemistry to that of the polymeric strips’ surface (i.e. surface wettability, surface energy and surface charge). We used Acrotest Pens to measure the surface tension and found that values below 42 mN/m and for layers thicknesses below 50 um remained uniform on the pre-treated PET strip without beading. The layer thickness was measured using the Keyence sensor LKG 150, as shown in Fig. S4.

A robotic arm by Epson moved the bio-chip between the SCT subsytem and imaging subsystem and the buffer bath for storage, as shown in Fig. 2b. Strips of 3M tape, 20 μm thick one cm wide, were stuck to the biochip and served as spacers, see Fig. S5. The industrial roller advanced the PET strip in a controlled manner to transfer the desired reagents to the DNBs attached to the biochip, which took 0.25 sec. for a roller speed of 80m/min.

To build the complementary DNA molecule for base calling we performed biochemical cycles similar to those for NGS flow cells. We were able to shorten the buffer exchange time from 1 minute to within 5 seconds for all steps, and thus one complete biochemical cycle took 1.5 minutes (including transfer time for robot).

After each biochemical cycle, the chip was transferred to the imaging subsystem of our DNBSEQ-T10 sequencer by the robot arm. Because of larger biochip, it was necessary to completely imaging all the DNBs which took less than one minute - see the video clip in supplementary information. The software from our DNBSEQ-T10 was used with no modification for base calling and other standard measurements such as lag and run on.

## Supporting information

video clip 1_SCT machine without cover

video clip 2_automation with T10

Simulation 1. 20um Gap_ buffer exchange ratio is 2X_18s for single volume pass

Simulation 2. one second is set for single volume pass

Simulation 3. buffer exchange is 5X__one second is set for single volume pass

## Conflicts of interest

The authors are employee of BGI-Shenzhen or MGItech.co and bound by confidentiality agreements.

## Acknowledgements

The work is supported by BGI-Shenzhen and MGItech.co., China Postdoctoral Science Foundation (2019M650215) and China National Gene Bank. We thank Chutian Xing for discussion and suggestion in engineering issues, Meihua Gong for instruction in biochemical process, Wangsheng Li for his help in biochip loading, Hong Xu for help in thickness determination, Ming Ni, Dong Wei for resource providing as well as Jian Liu for support of this project.

## Electronic Supplementary Information

**Fig. S1.**
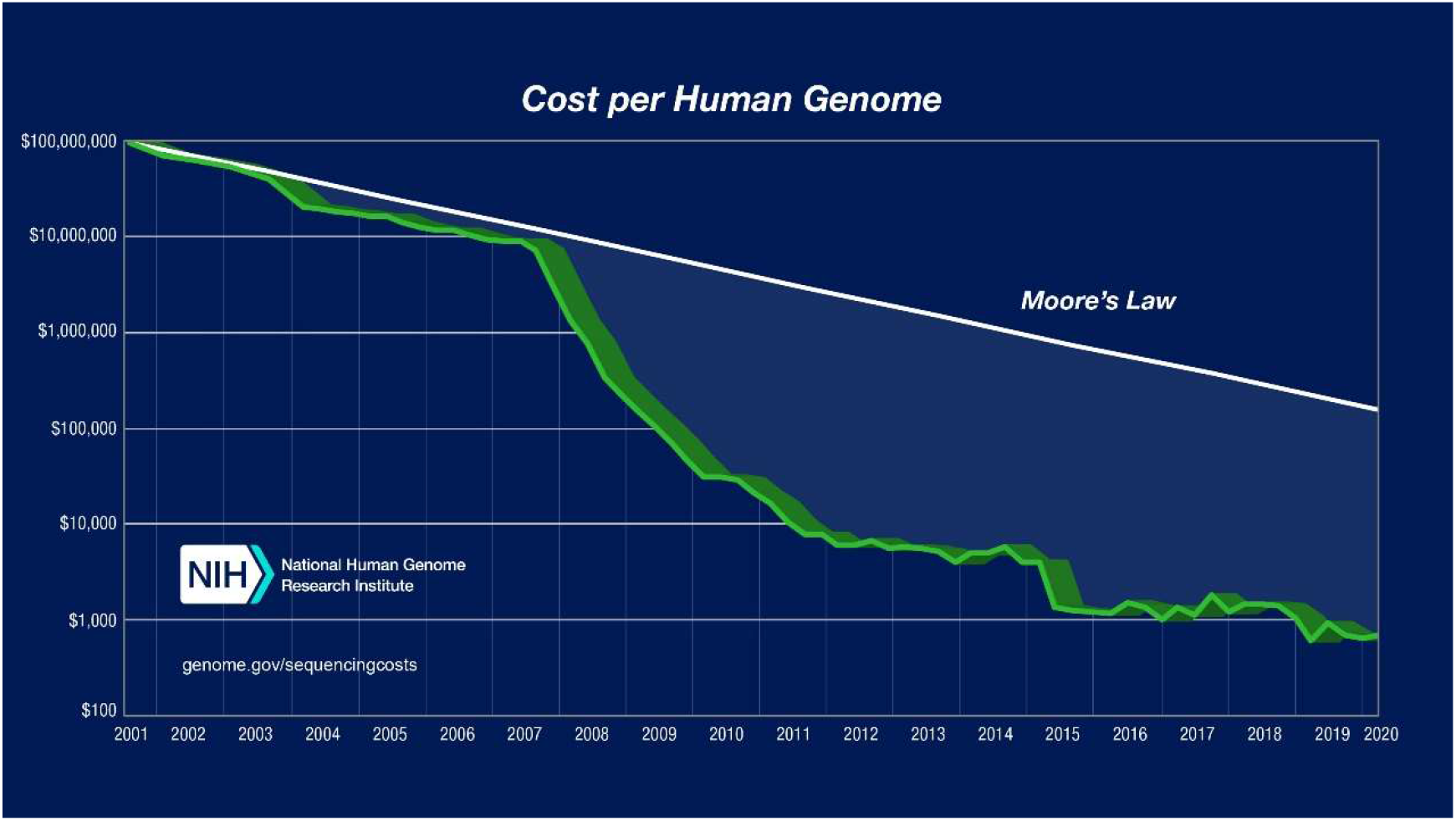
The trend of cost for whole genome sequencing (WGS), and the structure of a flow cell which is the well-established method for next-generation DNA sequencing (upper right corner). The next-generation sequencing (NGS) platform first released in the mid‑2000s heralded a 50,000 fold drop in the cost of human genome sequencing since the Human Genome Project^36^. For state-of-the-art flow cells technology, more than 80% of the consumable cost arises from reagents. Courtesy: National Human Genome Research Institute, https://www.genome.govabout-genomicsfact-sheetsDNA-Sequencing-Costs-Data. Illumina, Inc. https://www.illumina.com/company/news-center/multimedia-images.html.

**Fig. S2.**
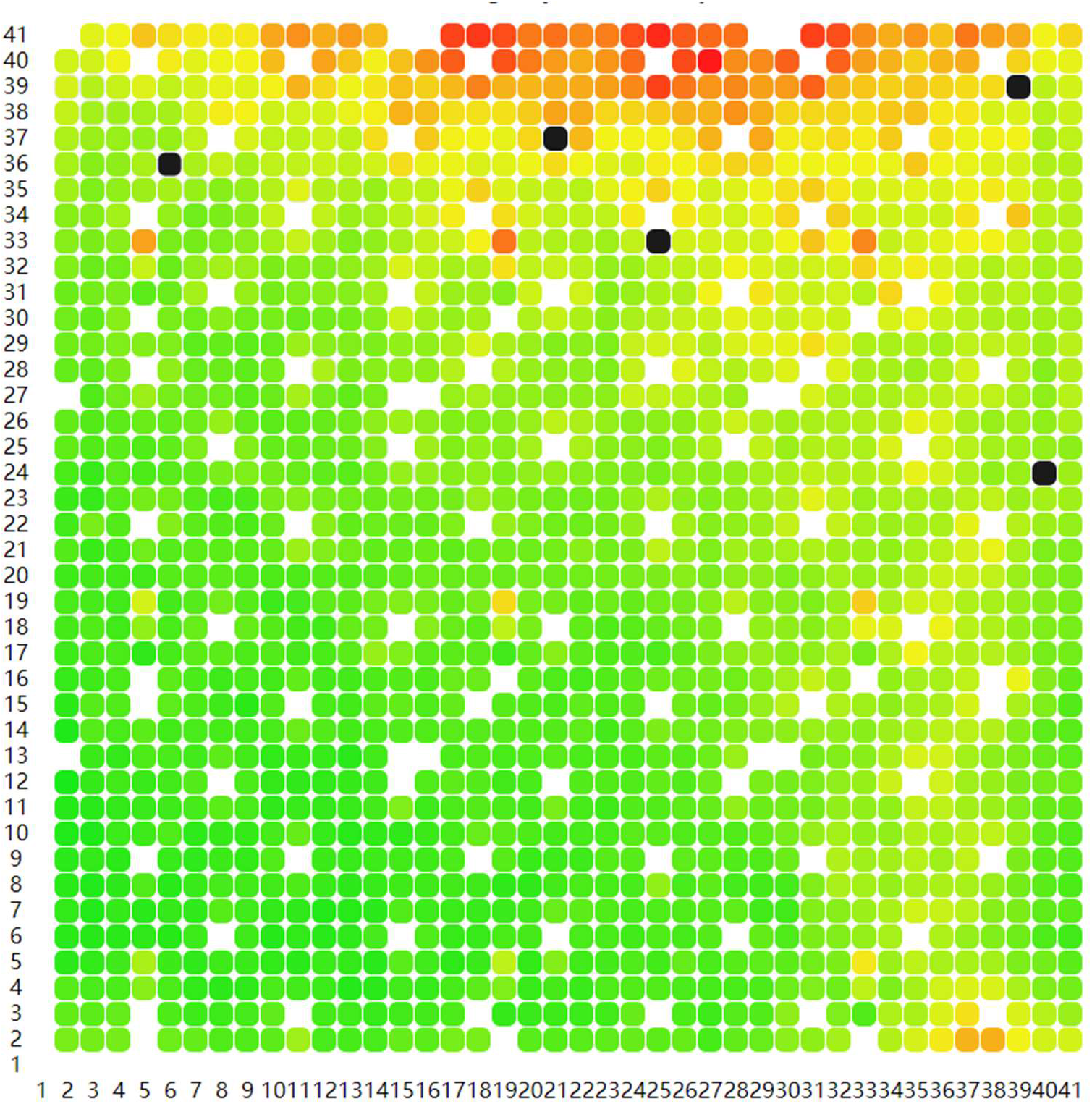
A BIC (basecalling information content) heatmap of a flow cell that experienced incomplete reagent replacement. Every red dot is a FOV (field of view) that evaluated as not good enough for basecalling.

**Fig. S3.**
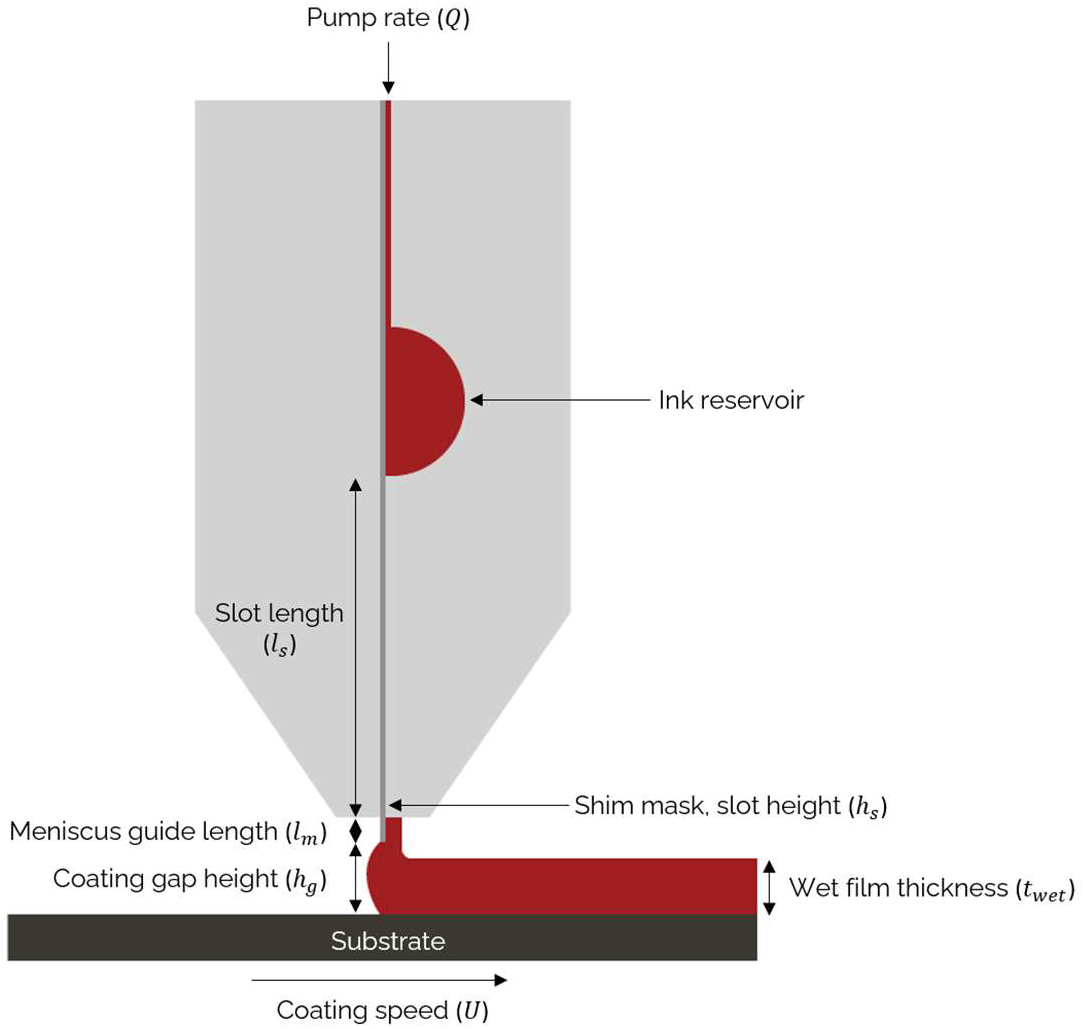
The schematic of a die for thin-layer reagent coating. Courtesy: FOM Technologies.

**Fig. S4.**
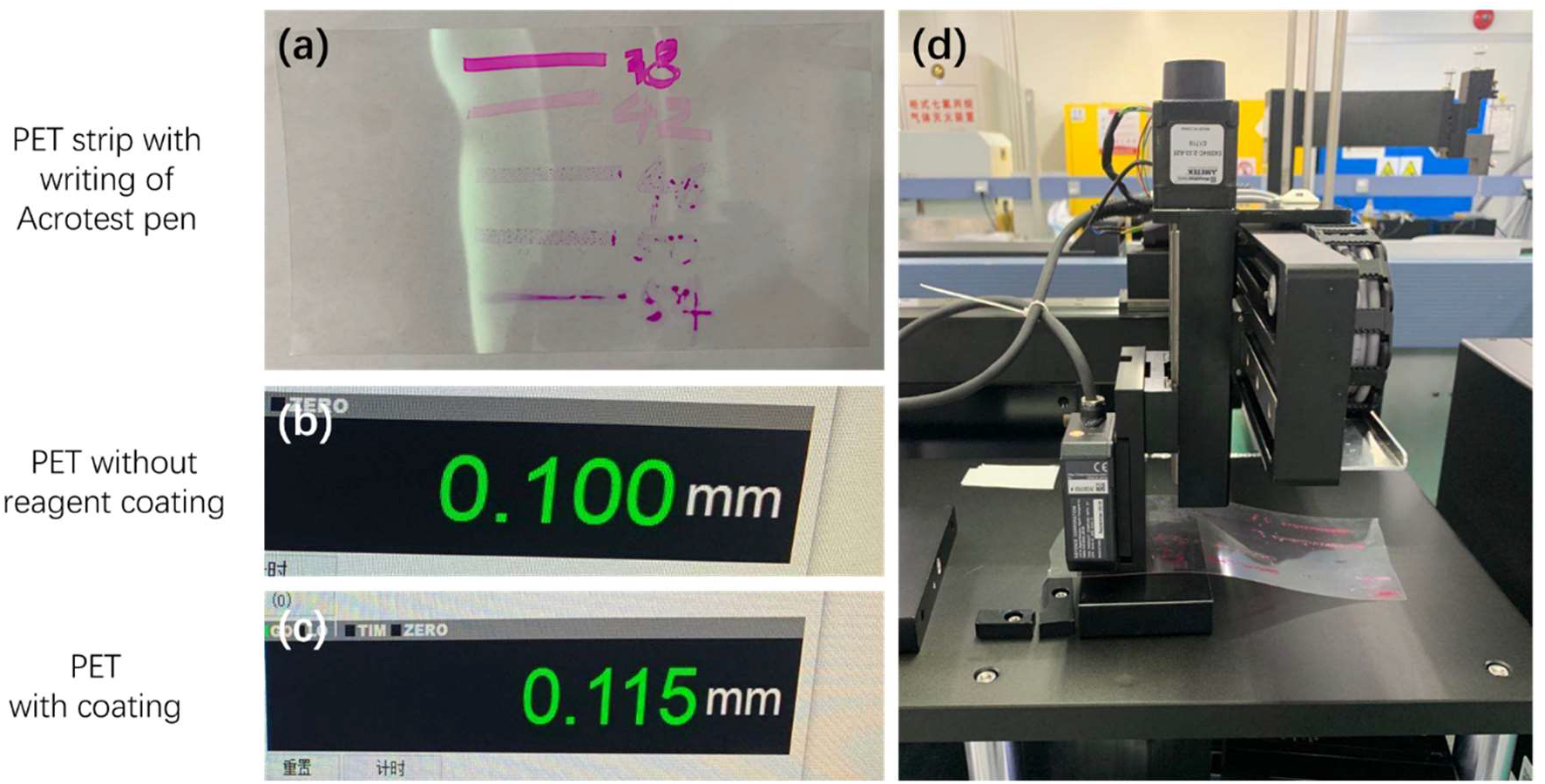
Uniformity and thickness test of reagent coating. (a) PET strip with writing by Acrotest pens to determine surface tension. Thickness display of PET membrane (100 μm) without (b) and with (c) reagent coating using (d) Thickness determination platform based on a Keyence sensor of LKG150.

**Fig. S5.**
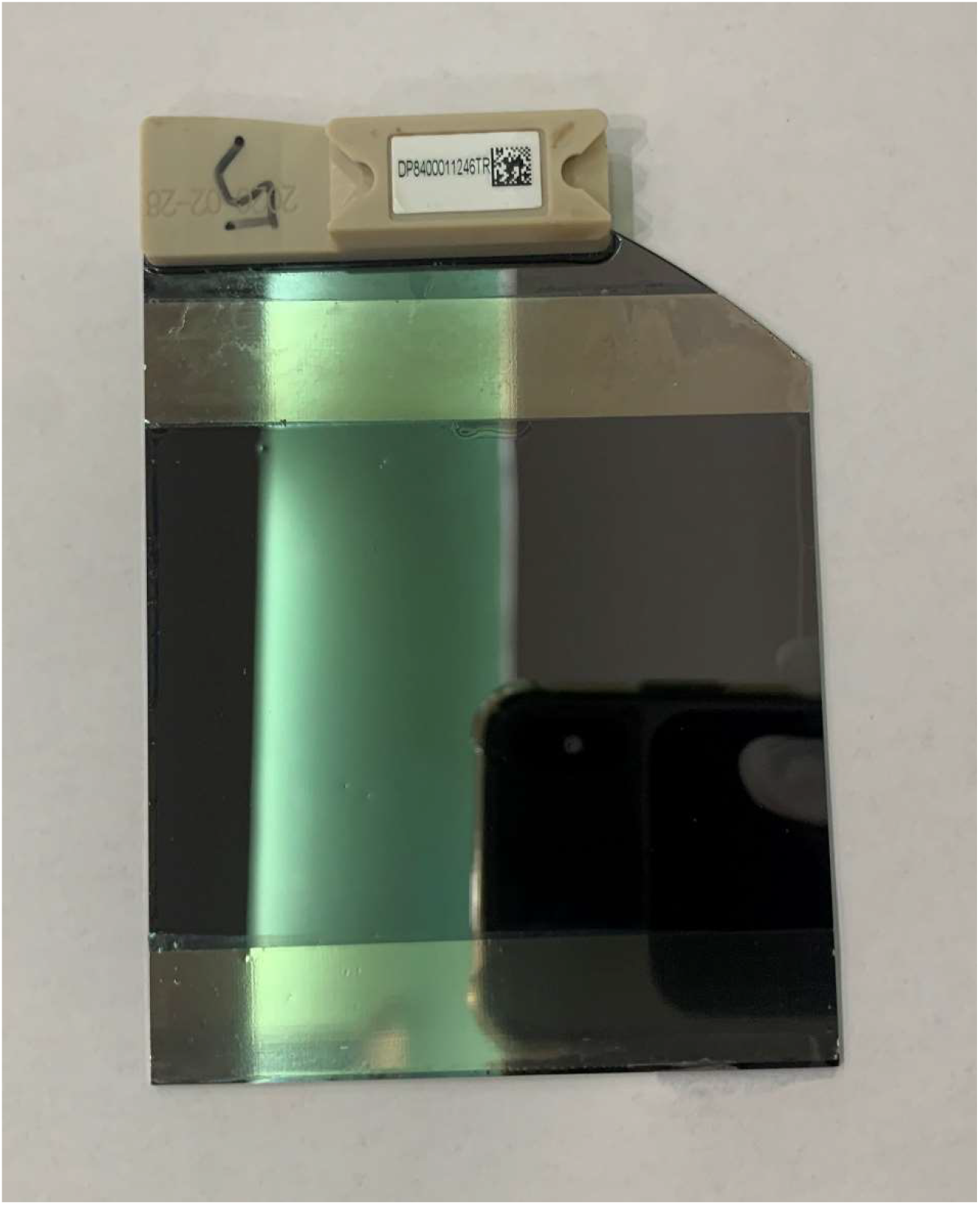
The images of a sequencing chip with two strips of 20 μm thick tape that act as spacers.

**Fig. S6.**
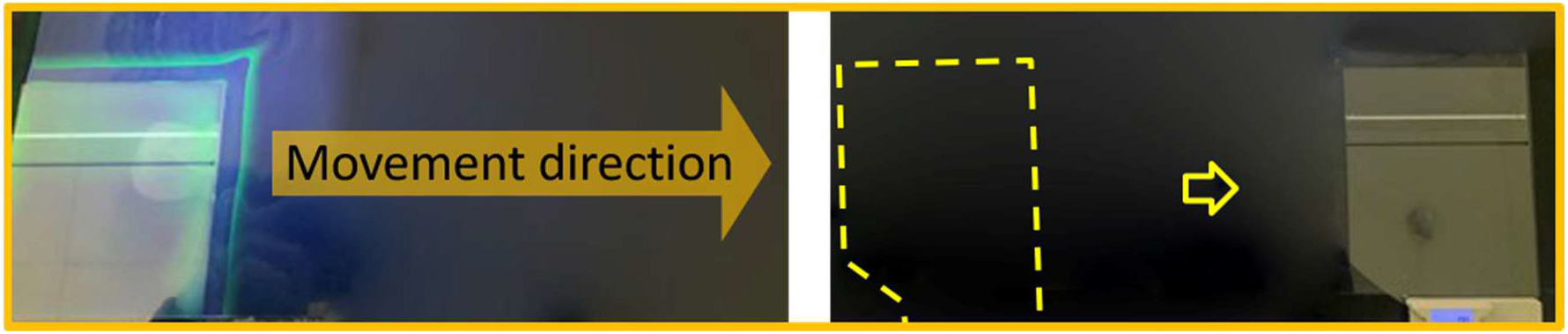
Replacing a fluorescent layer on the biochip with a buffer by advancing the PET strip.

## Calculation of reagent layer thickness for one NGS cycle

Expensive contents in NGS reagents include enzymes and dNTP (N stands for A, T, C, G). Since enzymes act as catalyst in biochemical reactions, we calculate dNTP consumption as follows:

The DNA nanoballs are patterned on the biochip to form a matrix with distance of 700 nm of two adjacent DNBs, assuming we have a 7 CM by 7 CM biochip, then:

Spot number = 7 CM × 7 CM ÷ (700 nm × 700 nm) = 10^10^

Assuming every DNB needs one fluorescent base (either A, C, T or G) in a cycle, and assuming consumption of A, C, T, G is the same for each cycle, and assuming each DNB needs ~1000 same fluorescent bases, then:

Consumption of bases of A, C, T or G = 10^10^ ÷ 4 × 1000 = 2.5×10^12^

For a normal bioreaction, the working concertation of dNTP is 0.5 μmol/L, then:

dNTP in 1 L reagent = 6.02 × 10^23^ × 0.5 × 10^−6^ = 3.01 × 10^17^
then total bases of A, C, T, or G in 1 L reagent = 3.01 × 10^17^ × 4 = 1.2 × 10^18^

For a 7 CM × 7 CM chip, dNTP reagent thickness of 1 L = 10 CM × 10 CM × 10 CM ÷ 7 CM × 7 CM = 20.4 CM = 2.04 × 10^5^ μm

So, the total number of bases in 1 μm thick dNTP reagent = 1.2 × 10^18^ ÷ (2.04 × 10^5^) = 5.88e^12^

Then, The required thickness of dNTP reagent = 2.5×10^12^ ÷ 5.88×10^12^ = 0.425×10^−3^ = 0.425 μm

The state-of-the-art flow-cell technique has a gap of 100 μm, assuming the buffer exchange rate is 5X, the actual reagent thickness of dNTP reagent = 100 μm × 5 = 500 μm.

The usage of reagent in state-of-the-art flow cell technology= 0. 425 μm ÷ 500 μm = 8.5×10^−4^ = 0.08 %

The reagent thickness of SCT is 20 μm and exchange ratio is 2X, then the usage of reagent is 0. 425 μm ÷ 20 μm × 2 = 1 %

Assuming 2 μm SCT is viable, then the usage of reagent is 0. 425 μm ÷ 2 μm × 2 = 10 %

## Notes

### Competing Interest Statement

The authors are employee of BGI-Shenzhen or MGI-Tech.co and bound by confidentiality agreements.

